# Maternal depression and child behaviours: sex-dependent mediation by glucocorticoid receptor gene methylation in a longitudinal study from pregnancy to age 5 years

**DOI:** 10.1101/187351

**Authors:** Jonathan Hill, Andrew Pickles, Nicola Wright, John P Quinn, Chris Murgatroyd, Helen Sharp

## Abstract

**Background:** Evolutionary hypotheses predict that male fetuses are more vulnerable to poor maternal conditions than females (Sex-biased Maternal Investment), but that the adaptive female fetus, with a more responsive hypothalamic-pituitary-adrenal (HPA) axis, is put at later risk of glucocorticoid mediated disorders where there is a mismatch between fetal and postnatal environments (Predictive Adaptive Response). Self-report measures of prenatal and postnatal depression and maternal report of child anxious depressed symptoms at 2.5, 3.5 and 5.0 years were obtained from an ‘extensive’ sample of first time mothers recruited from the general population (N = 794). Salivary *NR3C1* 1-F promoter methylation was assayed at 14 months in an ‘intensive’ subsample (N = 176) stratified during pregnancy by psychosocial risk. Generalised structural equation models (SEM) were fitted and estimated by maximum likelihood to allow inclusion of participants from both intensive and extensive samples.

**Results:** Postnatal depression was associated with *NR3C1* methylation and with anxious-depressed symptoms in the daughters of mothers lacking the hypothesised protective effect of high prenatal depression (prenatal-postnatal depression interaction for methylation, p =.00001; for child symptoms, p = .011). In girls, *NR3C1* methylation mediated the association between maternal depression and child anxious-depressed symptoms. The effects were greater in girls than boys, and the test of the sex differences in the effect of the prenatal-postnatal depression interaction on both outcomes gave χ^2^(2) = 5.95 (p=.051).

**Conclusions:** This is the first study to show in humans that, as a result of sex-biased reproductive investment and fetal adaptation, epigenetic and early behavioural outcomes may arise through different mechanisms in males and females. Epigenetic effects at the *NR3C1* promoter mediated mismatch between prenatal and postnatal maternal conditions and child anxious-depressed symptoms, specifically in females.

## BACKGROUND

The ‘fetal origins’ hypothesis was first proposed to account for associations between low birth weight and obesity, cardiovascular disease, and Type II diabetes in middle and old age [1]. According to this hypothesis, low birth weight reflects evolved adaptive mechanisms that confer advantages later in life in food scarce environments, but create risk in the presence of high calorie diets, common in industrialised societies. Far from being a mechanism specific to nutrition in humans, adaptations prior to birth that anticipate later environments are found across species, possibly reflecting a conserved ‘Predictive Adaptive Response’ (PAR) mechanism [2, 3]. According to the PAR hypothesis matched prenatal and postnatal conditions will be associated with good outcomes, while mismatching creates risks for later offspring adaptation. In relation to effects on psychiatric disorders, many studies have reported that anxiety, depression and behavioural symptoms in children are predicted by prenatal stressors, maternal depression and anxiety, and by low birth weight [4 - 9] however none has tested whether the PAR mechanism modifies these outcomes.

Fetal adaptations may additionally vary by sex of the fetus. According to the Trivers-Willard (T-W) hypothesis, if maternal health during pregnancy predicts later reproductive fitness in the offspring, then a male predominance of births will be favoured when maternal conditions are good, because healthy males compete successfully for females. By contrast, when maternal conditions are poor, the sex ratio will be reversed, both to avoid bearing males who compete less successfully for females, but also because, compared to females, health outcomes for mothers following male births are poorer [10]. Although this hypothesis has been subject to challenges and modifications [11], the central idea that reproductive strategies associated with poor maternal conditions involve sacrifice of males and protection of females has received substantial support. It is also consistent with well documented observations that male fetuses are more vulnerable to threats such as preterm birth, and are more likely to suffer neurodevelopmental consequences of fetal insults [12]. This hypothesis would appear to predict better outcomes for females following poor maternal conditions. However, if this protective effect in females arises from advantages conferred by fetal anticipation of matched environments (PAR hypothesis), mismatches between maternal conditions during pregnancy and the postnatal environment will create vulnerability. Combining the T-W and PAR hypotheses leads to the prediction that the effects of prenatal risks will operate differently in males and females. In females, vulnerability will be generated by particular combinations of prenatal and postnatal risks, while in males poor outcomes will arise incrementally from degree of prenatal risk. In the only human study we are aware of to have examined for the combined effects predicted by the T-W and PAR hypotheses, matched environments indexed by prenatal and postnatal depression (low-low and high-high) were associated with better cognitive and motor outcomes over the first year of life than mismatched prenatal and postnatal depression, and this effect was seen in females only [13]. However many studies have reported sex differences in developmental outcomes in relation to prenatal risks, without examining for the interplay with postnatal environments. Sex differences in fetal responses to stress [14], and in later emotional and behavioural problems following maternal anxiety or depression during pregnancy and low birth weight [4, 7, 8, 15, 16] have been identified.

In animal models, prenatal and postnatal stress cause long-term elevations in hypothalamic pituitary axis (HPA) reactivity and anxiety-like behaviours. This is mediated via reduced glucocorticoid receptor (GR) gene *NR3C1* expression, particularly in the hippocampus which impairs HPA axis feedback mechanisms [17]. The epigenetic process of DNA methylation involves the addition of methyl groups to CpG dinucleotides in gene regulatory regions that are associated with repressed gene expression. Animal findings of the epigenetic effects of early life stress have been translated to humans in a study reporting elevated *NR3C1* 1-F promoter methylation and reduced *NR3C1* expression in post-mortem hippocampal tissue of suicide completers who were abused during childhood, when compared to non-abused [18]. Other studies using peripheral DNA from blood or saliva of infants and adolescents have shown increased levels of *NR3C1* methylation associated with prenatal and childhood adversities [19, 20, 21]. Several clinical studies examining leukocytes have reported elevated methylation of the homologous human *NR3C1* 1-F promoter (homologous to the rat 1-7 promoter) at a specific CpG (CpG unit 22,23, Figure 1) associated with prenatal maternal depression [19, 22-24] and childhood stress [25]. Studies in humans also find associations between prenatal anxiety and postnatal depression in mothers, and adolescent depressive symptoms mediated via HPA axis dysregulation [26, 27], consistent with the role of HPA axis dysregulation in adolescent depression [28]. Higher *NR3C1* methylation levels, hypothesised to contribute to reduced *NR3C1* expression (18), have been associated with increased salivary cortisol stress responses in infants at 3 months [19] and a flattened cortisol recovery slope following stress in adolescents [29], suggesting methylation of *NR3C1* may impair negative feedback of the HPA axis.

**Figure 1.**
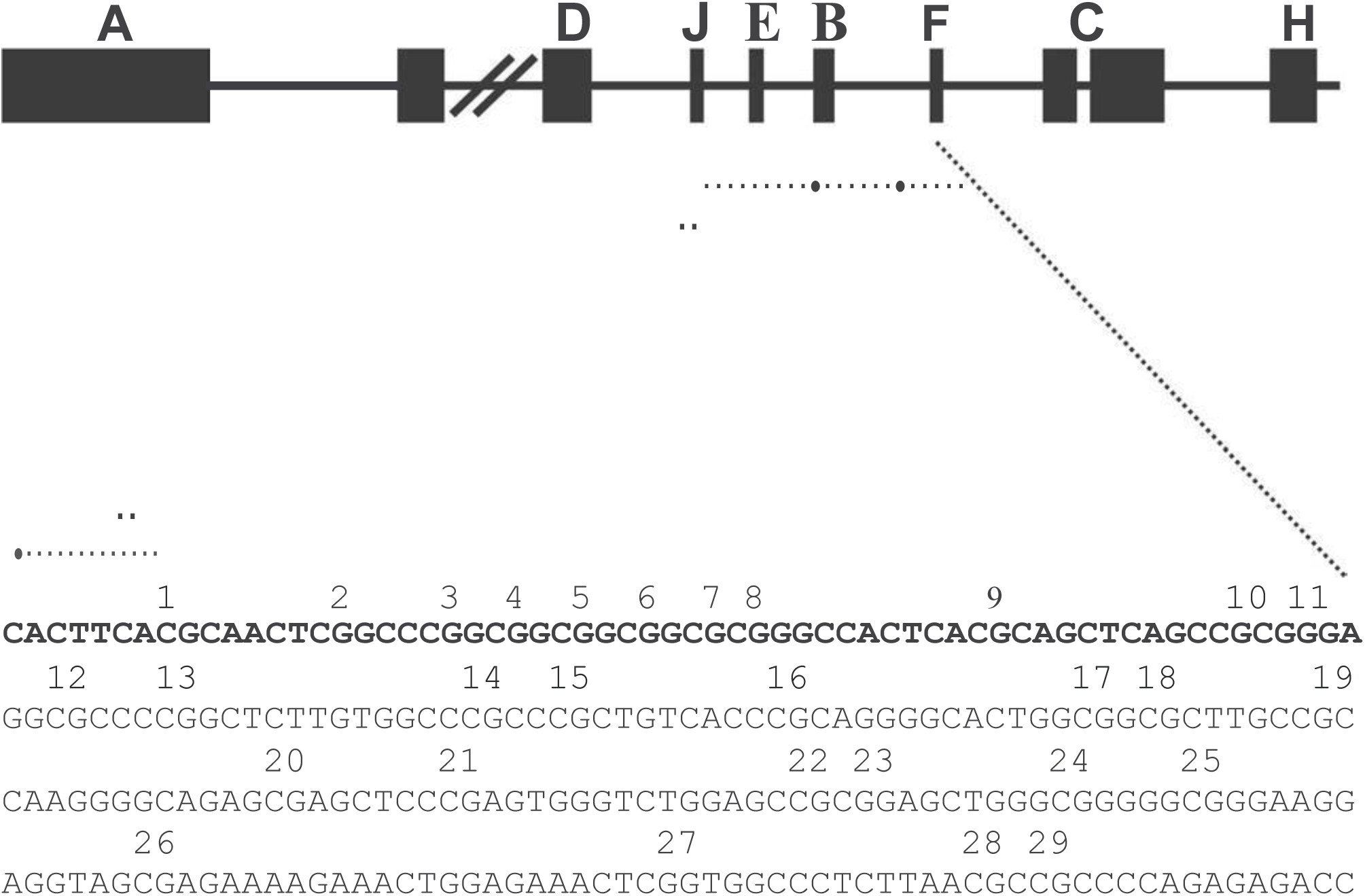
Scheme of the human NR3C1 gene analyzed by bisulfite pyrosequencing. The 5′-end of the human NR3C1 gene contains multiple first exons, with multiple transcriptional start sites and mRNA splice variants. The region analyzed by bisulfite pyrosequencing (primer sequences are in bold) contains 29 CpGs and encompasses exon 1-F, which is the human homolog of the rat exon 1–7, previously shown to be differentially methylated [47]

In the first study to examine the interplay between prenatal and postnatal depression in relation to *NR3C1* gene methylation, we showed that the association between postnatal maternal depression and *NR3C1* 1-F promoter methylation in their children was stronger where mothers had reported lower depression during pregnancy, in line with the PAR hypothesis [30]. However, we did not examine for sex differences. Sex differences in glucocorticoid mechanisms associated with prenatal stress have been shown in animal models. In rats many effects of prenatal stress on later development are seen only in females, and these are abolished by adrenalectomy [31]. The effects predicted by a combination of the T-W and PAR hypotheses, have been demonstrated in starlings where mismatched prehatch-posthatch conditions had a greater effect on corticosterone levels in female than male chicks, but prenatal risk increased mortality in male chicks [32, 33]. In humans, a sex difference in associations between prenatal depression and *NR3C1* 1-F promoter methylation has been reported [34], although the interplay with postnatal depression however was not analysed.

In this study we examined predictions based on the T-W sex-biased parental investment and PAR hypotheses. In females, where individual and species vulnerability are reduced by matching environments but increased by mismatching, the presence of good prenatal conditions followed by adverse rearing experiences, and *vice versa*, will create vulnerability to child anxiety and depression. Based on the animal models, we predicted this effect in females will involve altered HPA axis reactivity arising from epigenetic modifications of the GR gene. In males, where individuals are sacrificed for species advantage, the presence of prenatal stress will create vulnerability, unmodified by later environment quality. The animal models suggest that glucocorticoid mechanisms are implicated in excess male deaths under unfavourable maternal conditions, but they may not contribute to effects of prenatal stress on functioning after birth.

These predictions were tested in a longitudinal study using measures of prenatal and postnatal depression, of *NR3C1* 1-F promoter region methylation at 14 months of age, and anxious depressed symptoms in children across the preschool period. We predicted that in girls but not boys, low prenatal depression followed by high postnatal maternal depression, and high prenatal depression followed by low postnatal depression will be associated with elevated anxious depressed symptoms and elevated *NR3C1* methylation. In boys, prenatal and postnatal depression will be independent risks for elevated anxious-depressed symptoms, without the interaction between them predicted for females.

## METHODS

### Design

The participants were members of the Wirral Child Health and Development Study, a prospective epidemiological longitudinal cohort of first-time mothers recruited in pregnancy to study prenatal and infancy origins of emotional and behavioural disorders. The full cohort of 1233 mothers with live singleton births have participated in several waves of assessment with a stratified random sub-sample of 316 identified for additional, more intensive assessment (intensive sample). Strata were defined on the basis of low, medium and high psychosocial risk (scores of <=2, 3 or >3 on an inter-partner psychological abuse scale provided on entry to the study at 20 weeks of pregnancy), with higher selection probabilities for those at higher risk. Appropriately analysed, the design allows estimates of means and coefficients for the whole general population cohort to be derived even for measures available only in the intensive sample [35].

Approval for the procedures was obtained from the Cheshire North and West Research Ethics Committee (UK) (reference no. 05/Q1506/107). The extensive sample was identified from consecutive first time mothers who booked for antenatal care at 12 weeks gestation between 12/02/2007 and 29/10/2008. The booking clinic was administered by the Wirral University Teaching Hospital which was the sole provider of universal prenatal care on the Wirral Peninsula. Socioeconomic conditions on the Wirral range between the deprived inner city and affluent suburbs, but with few from ethnic minorities. The study was introduced to the women by clinic midwives who asked for their agreement to be approached by study research midwives when they attended for ultrasound scanning at 20 weeks gestation. After complete description of the study to the women, written informed consent was obtained by the study midwives, who then administered questionnaires and an interview in the clinic.

### Participants

Of those approached by study midwives, 68.4% gave consent and completed the measures, yielding an extensive sample of 1233 mothers with surviving singleton babies. The sampling flow chart has been published previously [35]. The mean age at recruitment of extensive sample participants was 26.8 years (s.d.5.8, range 18-51). Using the UK Index of Multiple Deprivation (IMD) [36] based on data collected from the UK Census in 2001, 36.6 % of the extensive sample reported socioeconomic profiles found in the most deprived UK quintile, consistent with the high levels of deprivation in some parts of the Wirral. Forty eight women (3.9%) described themselves as other than White British.

In addition to assessments of the mothers at 20 weeks gestation, mothers and infants provided data at birth and postnatally at 5, 9, and 29 weeks, and at 14.19, s.d. 1.71 months (‘14 months’), 30.86, s.d. 2.31 months (‘2.5 years’), 41.90 s.d. 2.48 months (‘3.5 years’) and 58.64 s.d. 3.74 months (‘5 years’). Two hundred and sixty eight mothers and infants came into the lab at 14 months for detailed observational, interview and physiological measures. This was the first occasion at which saliva for DNA was collected. Seven parents declined consent for DNA collection, 3 samples were spoilt, and 25 assessments were curtailed before saliva collection because of time constraints. Sufficient DNA for methylation analyses was obtained from 181 infants. Maternal reports of child anxious-depressed symptoms were available on 253 of the intensive sub-cohort at 2.5 years, on 825 of the whole cohort at 3.5 years and on 768 of the whole cohort at 4.5 years.

### Measures

#### Maternal depression

Maternal symptoms of depression were assessed at 20 weeks gestation and at every follow up point using the Edinburgh Postnatal Depression Scale (EPDS), which has been used extensively to assess prenatal and postnatal depression [37].

#### DNA methylation

Methylation status in the *NR3C1* 1-F promoter was examined at the same CpGs (CpG unit 22 and 23, shown in Figure 1) identified in previous studies (24). DNA collected from Oragene® saliva samples, was extracted, bisulphite treated, amplified (Forward, GACCTGGTCTCTCTGGGG; Reverse, TGCAACCCCGTAGCCCCTTTC) and run on a Sequenom EpiTYPER system (Sequenom Inc., San Diego, US), providing an average for methylation across the two CpG units. Data was transformed to percentage of methylation at CpG unit 22 and 23 to allow for comparison with previous analysis of differential methylation at this locus.

#### Child anxious-depressed symptoms

Child symptoms were assessed by maternal report at 2.5, 3.5 and 5.0 years using the preschool Child Behavior Checklist (CBCL) [38]. It has 99 items each scored 0 (not true), 1 (somewhat or sometimes true), and 2 (very true or often true), which are summed to create seven syndrome scales. Only the anxious/depressed scale was analysed for this report, and as recommended in the CBCL manual, raw scores were used [39].

#### Stratification variable and confounders

Partner psychological abuse was assessed using a 20 item questionnaire covering humiliating, demeaning or threatening utterances in the partner relationship during pregnancy over the previous year [40]. Maternal age (at this first pregnancy), marital status at 20 weeks gestation, and socioeconomic status were included as covariates because of their established associations with adult depression. Socioeconomic status was determined using the revised English Index of Multiple Deprivation (IMD) [36] based on neighborhood deprivation. All mothers were given IMD ranks according to the postcode of the area where they lived and assigned to a quintile, based on the UK distribution of deprivation. Mother’s years of education at enrolment in the study was recorded. Information about smoking was obtained at 20 and 32 weeks gestation and was included because of published associations with altered DNA methylation [41]. Birth records provided sex of infant, one-minute Apgar score, and birth weight and gestational age, from which a measure of fetal growth was obtained. Low fetal growth is associated with elevated fetal glucocorticoid exposure and so might be associated with elevated *NR3C1* gene methylation. Obstetric risk was rated using a weighted severity scale developed by a collaboration of American and Danish obstetricians and paediatric neurologists [42].

### Statistical Analysis

All analyses were undertaken in Stata 14 (StataCorp, 2015). Generalised structural equation models (SEM) were fitted using the sem procedure and estimated by maximum likelihood to allow inclusion of participants from both intensive and extensive strata. The anxiety-depression scores at 2.5, 3.5 and 5.0 years and *NR3C1* percent methylation at 14 months were highly skewed so scores were log-transformed and Winsorized at 2.5 standard deviations to reduce their skew. For further robustness, we report standard errors and p-values based on the heteroscedastic consistent estimator of the parameter covariance matrix. The main analyses included the stratification variable and confounds except for perinatal confounds as they may lie on a mediational pathway from prenatal depression, however the effect of adding those variables was examined. Model estimates and tests allowed for differential missingness associated with any of the covariates and observed responses included in the model, accounting for the stratified study design.

The pre-post environment mismatch predictions on both methylation and child symptoms were examined first by testing for two-way interactions between prenatal and postnatal depression in models estimated separately in females and males. We then tested for the sex difference by examining the three-way, sex by prenatal depression by postnatal depression interactions in a model that included both genders. The effects of combinations of prenatal and postnatal depression giving rise to these interaction effects are shown in the figures. The prediction of additive effects of prenatal and postnatal depression in boys was examined in models without interaction terms.

In the fitted models methylation was specified as a factor, measured without error by the observed methylation, a device that implicitly imputes rates of methylation where these have not been observed, but doing so in a manner which recognises our uncertainty in these unobserved values. This enables participants with partial data that would be informative for some parts of the model to be included.

## RESULTS

### Descriptive Statisticsd

Table 1 gives summary statistics for males and females separately for the measures included in the analysis, and shows the sample size at each data collection point. As described in the statistical analysis section, differences in the available sample for different measures were accounted for by use of weighted, maximum likelihood or covariate adjusted estimators. Figure 2 shows the structure of the SEM model in which maternal history of depression predicts *NR3C1* methylation (solid red arrows) and maternal report of child anxious-depressed symptoms (solid black arrows). These analyses included the 412 girls and 382 boys on whom there were measures of maternal depression and maternal report of child anxious-depressed symptoms at a minimum of one follow up point as well as all confounders.

**Table 1.**
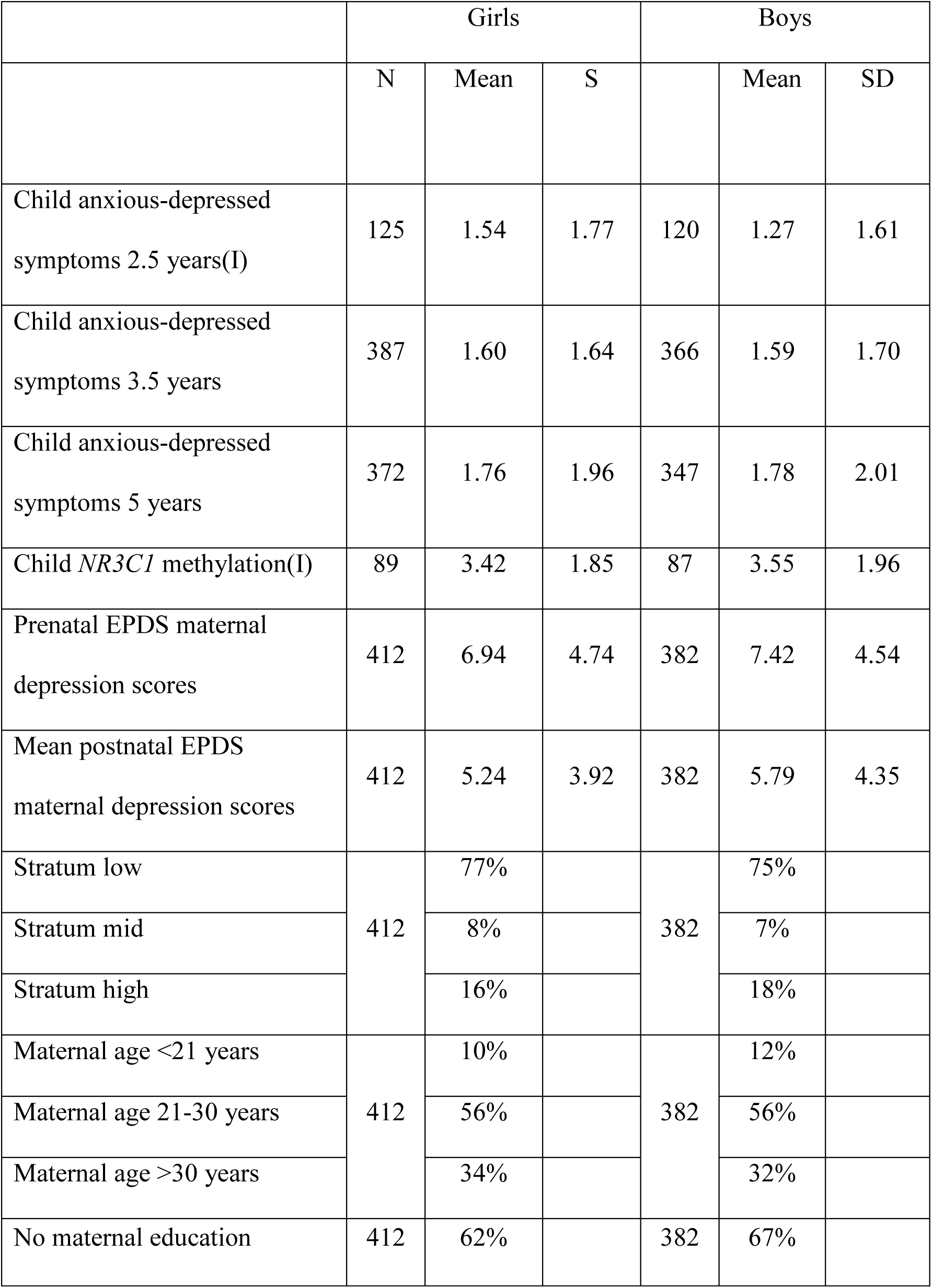
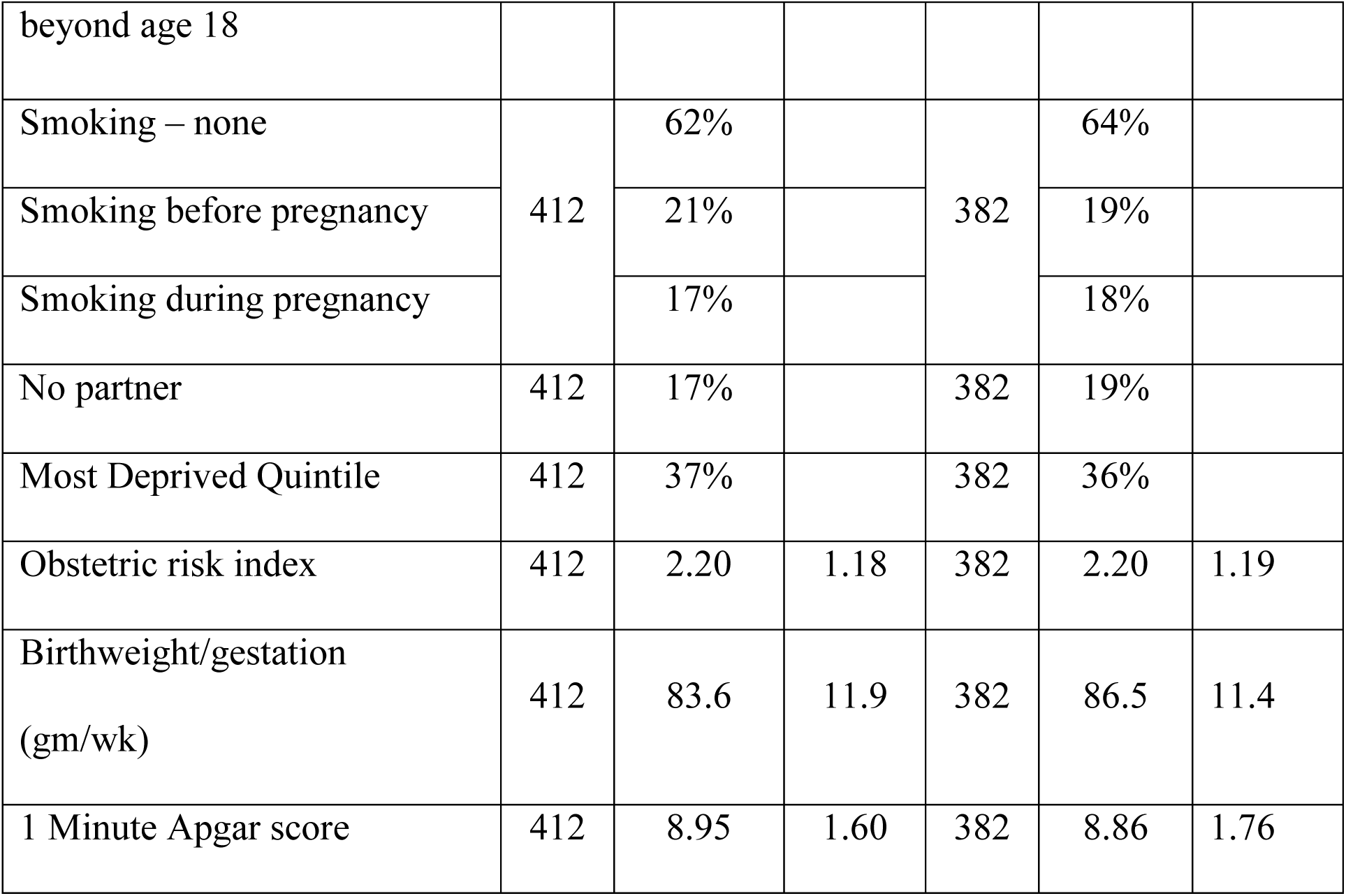
Summary statistics for outcomes, predictors and variables included as potential confounders for the modelled sample (I = measure based on intensively assessed sub-sample only)

**Figure 2.**
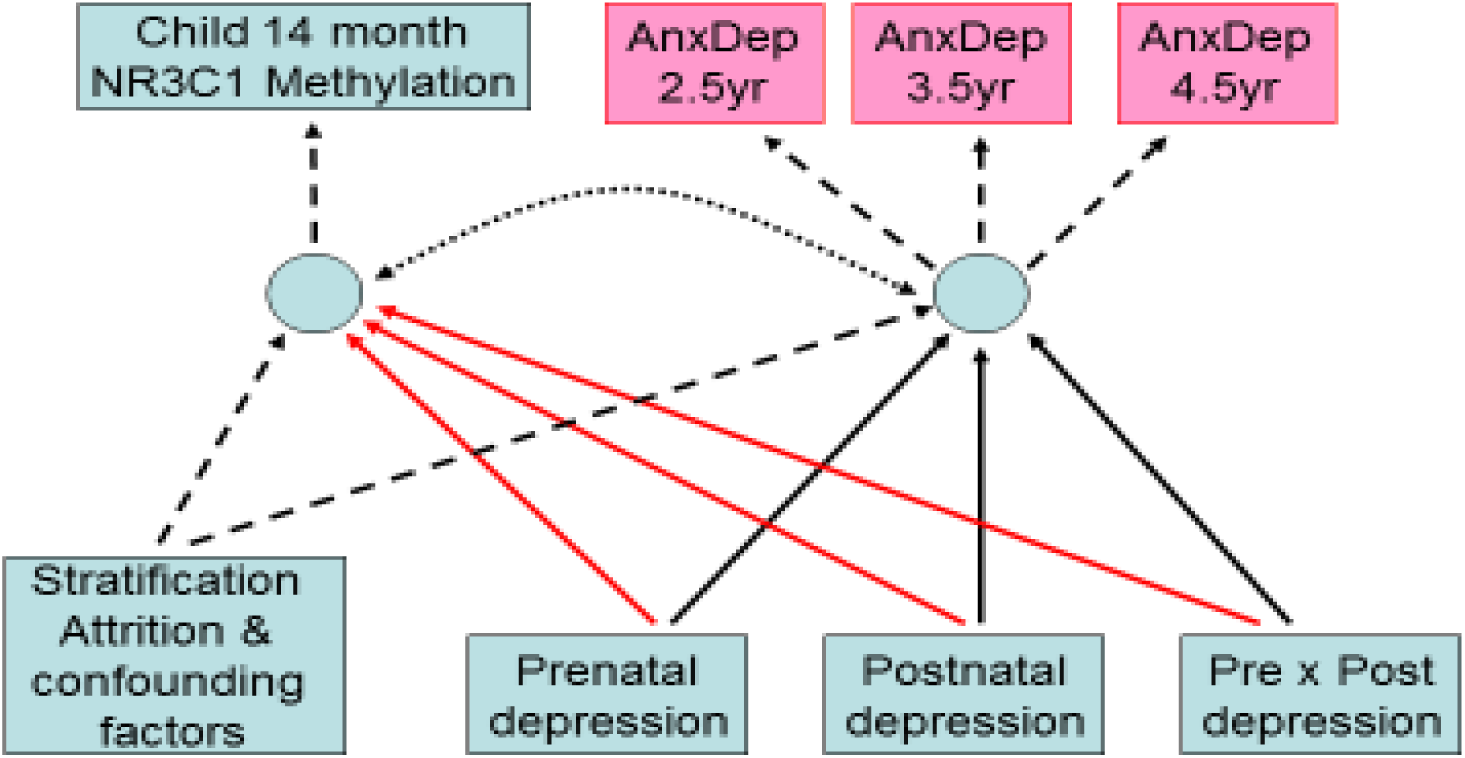
Structural equation model fitted to *NR3C1* 1-F promoter methylation at 14 months and CBCL anxious-depressed scores at ages 2.5, 3.5, and 5 years

Table 2 shows for girls and boys separately the estimated path coefficients from the standardised prenatal depression, postnatal depression and their interaction (product) of primary interest accounting for the stratification, attrition and confounders. We first tested the prediction that there would be an interaction between prenatal and postnatal depression in girls but not in boys. In girls there was a significant effect of the interaction between prenatal and postnatal depression on both child anxiety-depression (p=.011) and *NR3C1* 1-F promoter methylation (p =.00001). For boys, by contrast, anxious-depressed symptoms were not predicted by the prenatal and postnatal depression interaction term (p=.920), and the effect on *NR3C1* methylation was smaller than for girls, though still significant (p=.003). Adding the three additional potential confounders that were assessed after the prenatal exposure (obstetric risk index, 1-minute Apgar score and birthweight/gestational age) made no material difference to these associations. Fitting this model to boys and girls together, but allowing the effects of prenatal and postnatal depression exposure on the two correlated outcomes to differ by sex (in addition to a gender main effect), a Wald test of the sex differences in the effect of the prenatal-postnatal depression interaction on both outcomes (a difference of 0.20 for anxiety-depression and 0.18 for methylation) gave χ^2^(2) of 5.95 (p=.051), with the two individual interactions contributing equally (anxiety-depression p=.088, methylation p=.069).

**Table 2.** Summary of SEM analyses predicting *NR3C1* 1-F promoter methylation and child anxious depressed symptoms The table shows standardized factor loadings of child CBCL anxious-depressed symptoms at ages 2.5, 3.5 and 5 years, and main effects and effects of interaction of prenatal and postnatal depression in the prediction of the anxious-depressed factor and the *NR3C1* 1-F promoter methylation (effects of stratification factors and confounders not shown). Anxious-depressed symptoms and methylation are analysed together as correlated outcomes in an SEM. Coefficients for the effects of confounders and stratification factors are not shown (stratum, maternal age, maternal smoking, maternal education, no partner, neighbourhood deprivation).

|  | Female (n=412) |  | Male (n=382) |  |
| --- | --- | --- | --- | --- |
|  | Std Coeff | 95% C.I | Std Coeff | 95% C.I |
|  | [p-value] |  | [p-value] |  |
| Effects on child symptoms of anxiety-depression |  |  |  |  |
| Prenatal maternal depression | -0.06 | -0.23, 0.11 | 0.16 | -0.00, 0.33 |
| Postnatal maternal depression | 0.21 | 0.05, 0.38 | 0.17 | 0.03, 0.31 |
| Prenatal-postnatal interaction | -0.19<br>[p=.011] | -0.34,-0.05 | 0.01<br>[p=.920] | -0.11, 0.12 |
| Effects on child *NR3C1* 1-F promoter methylation |  |  |  |  |
| Prenatal maternal depression | -.02 | -0.28, 0.24 | -0.11 | -0.34, 0.12 |
| Postnatal maternal depression | 0.45 | 0.16, 0.75 | 0.38 | 0.11, 0.65 |
| Prenatal-postnatal interaction | -0.39<br>[p=.00001] | -0.56, -0.21 | -0.21<br>[p=.003] | -0.32, -0.08 |
| Child anxious-depressed symptoms factor loadings |  |  |  |  |
| 2.5 years | 0.81 |  | 0.72 |  |
| 3.5 years | 0.80 |  | 0.67 |  |
| 5 years | 0.57 |  | 0.81 |  |

We then tested the prediction that in boys there would be independent and additive effects of prenatal and postnatal depression, by estimating the model (not shown in the Table) for boys without the interaction term. This showed a significant effect on child anxiety-depression of postnatal depression (standardised coefficient 0.17, CI 0.04 to 0.30, p = .011) and an effect of similar magnitude, that was non-significant, of prenatal depression (0.15, CI - 0.02 to 0.33, p=.080). Independent effects on methylation were not seen (prenatal 0.05, CI - 0.17 to 0.27, p=.640; postnatal 0.13, CI -.09 to 0.36, p=.241).

Figure 3 displays how the interactions between prenatal and postnatal depression in the prediction of anxious-depressed symptoms differed between girls and boys. It can be seen that, in girls, at a low level of prenatal depression (1 standard deviation below the mean), increasing postnatal depression was strongly associated with increasing child anxious-depressed symptoms, while at a high level there was no association. With prenatal depression at the mean, the association was intermediate between the low and high prenatal levels. In boys, by contrast, as evidenced in parallel regression lines, there was no interplay between prenatal and postnatal maternal depression.

**Figure 3.**
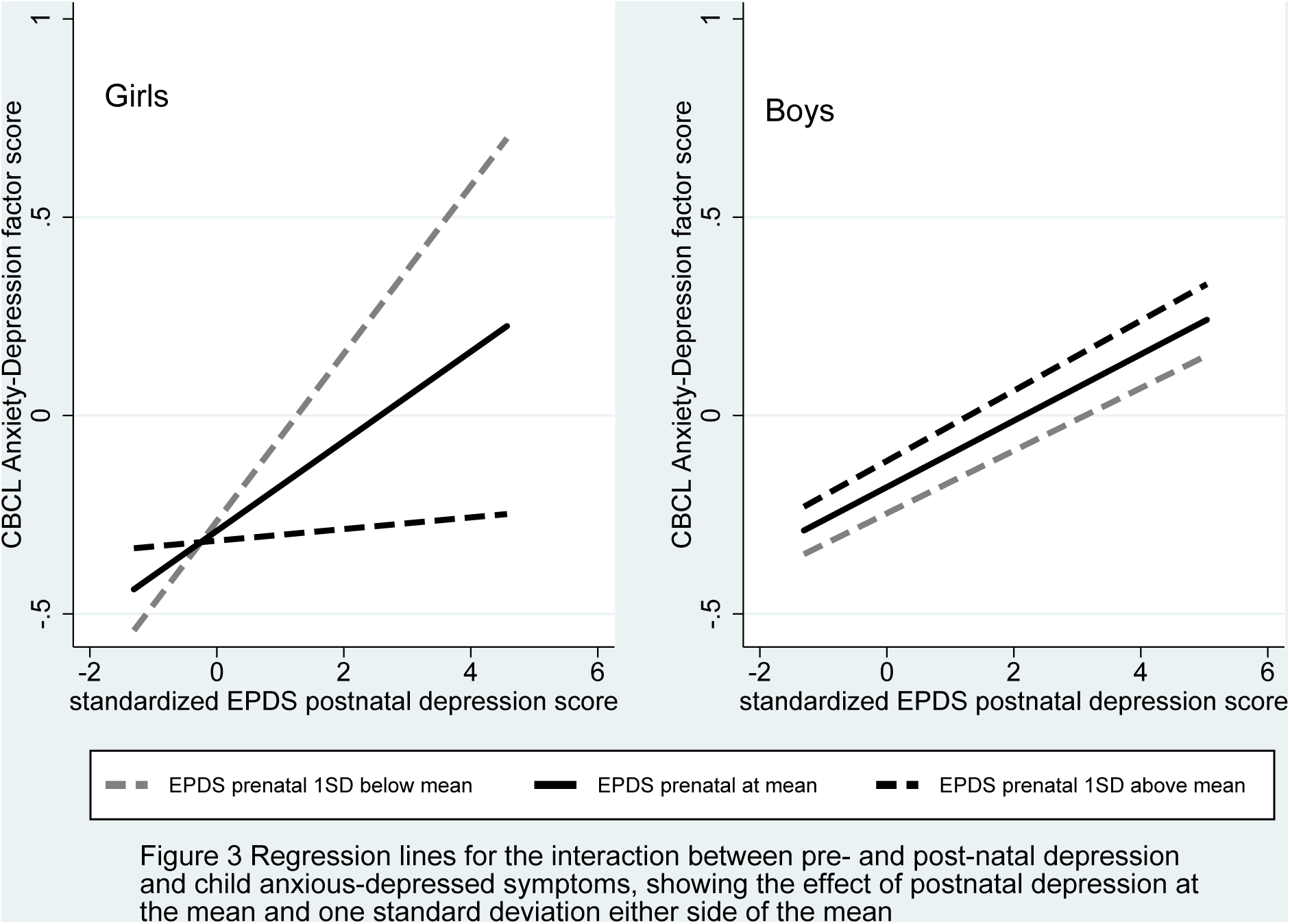
Regression lines for the interaction between pre- and post-natal depression and child anxious-depressed symptoms, showing the effect of postnatal depression at the mean and one standard deviation either side of the mean

As shown in Figure 4, the effects of prenatal-postnatal mismatch on methylation were again strongly evident in girls, with the greatest association between postnatal depression and methylation in the presence of low prenatal depression, and progressively weaker associations at higher levels of prenatal depression. The progressive effect of prenatal depression was also evident in boys but it was less strong.

**Figure 4.**
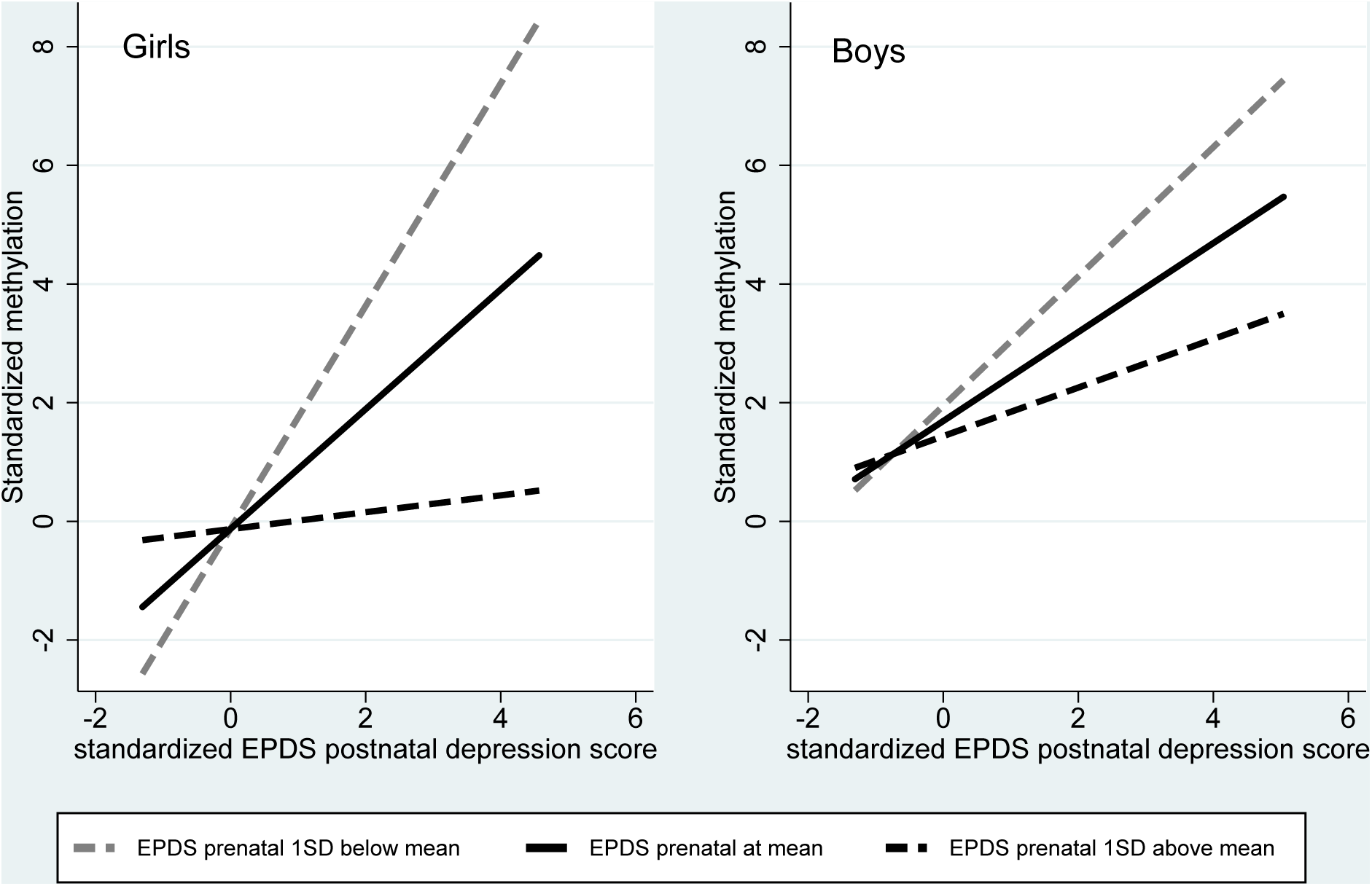
Regression lines for the interaction between pre- and post-natal depression and child NR3C1 1-F promoter methylation, showing the effect of postnatal depression at the mean and one standard deviation either side of the mean

In girls, replacing the correlation between the methylation and anxiety-depression factors by a causal effect, higher *NR3C1* methylation at 14 months was associated with higher anxiety-depressed symptoms (standardised coefficient 0.36 CI 0.05 to 0.67, p=.025). The residual direct effect of the prenatal-postnatal interaction on child anxiety-depression symptoms was substantially reduced, from -0.19 (shown in Table 2) to -0.06 (CI -0.26 to 0.15), becoming wholly nonsignificant (p=.600). For boys there was no evidence of an effect of methylation on symptoms (standardised coefficient -0.03, CI -0.31 to 0.24, p=.820).

## DISCUSSION

Many, although not all, of our predictions based on the evolutionary T-W and PAR hypotheses for sex-biased parental investment and fetal programming were supported in this longitudinal study, from 20 weeks of pregnancy and over the first 5 years of children’s lives. Mismatching between prenatal and postnatal maternal depression was associated with greater anxious-depressed symptoms and *NR3C1* methylation in girls. Both effects were most evident in girls exposed to high levels of postnatal depression. Their symptoms and *NR3C1* methylation were higher where their mothers had reported low levels of depression during pregnancy, in line with the idea that they had not been prepared by the fetal environment for postnatal exposure to maternal depression. In girls only, elevated *NR3C1* was associated with higher anxious-depressed symptoms, and mediated the association between maternal depression and child symptoms. In boys there was no evidence of effects of prenatal – postnatal mismatch on anxious depressed symptoms. However, and contrary to our prediction, the prenatal-postnatal mismatch effect on *NR3C1* methylation was seen in boys as well as in girls, although the size of the effect was smaller.

The strengths of the investigation include prospective study with a general population sample, accounting for a number of plausible confounds and factors associated with attrition. Also, by using SEM to create a latent variable from measurement at 3 time points over 2.5 years we reduced the risks arising from multiple testing for each time point, and we were able to examine the predictions in relation to persistently elevated symptoms likely to confer risk for an elevated trajectory for anxious-depressed symptoms over childhood [43]. The method adopted for missing methylation data exploited the properties of maximum likelihood for accounting for data assumed missing at random. Most missingness was by design because of the systematic stratification of the intensive sample, thus meeting this assumption, and inclusion of multiple covariates allowed us account for unplanned attrition. It is nevertheless possible that not all the necessary confounds to deal with non-random missingness were identified.

There were four principal limitations in relation to the measurement of *NR3C1* methylation. First, peripheral cell samples, both from blood and saliva, are heterogeneous, which may account for some of the variability in methylation. This can introduce a confound where other variables are associated with cellular heterogeneity [44]. Second, while studies combining peripheral cell and CNS post mortem estimations suggest that they are often substantially correlated [45], it cannot be assumed that DNA methylation in peripheral tissues reflects methylation in relevant CNS regions. This is particularly a concern because of substantial variations in epigenetic effects across brain regions and cell types. Third there are many combinations of CpG sites, even on a relatively circumscribed region such as the *NR3C1* 1-F promoter that could be examined, leading to the risk of multiple analyses and ‘significant’ findings occurring by chance. Fourth, although we accounted for a number of plausible confounds, environmental variables other than those included in analyses may better account for the findings.

No one study can establish the validity of estimates of peripheral cell methylation as indices of CNS methylation, however a finding of the same pattern of associations for peripheral cell methylation and for behaviours that undoubtedly reflect CNS function, and for mediation of the association between maternal depression and symptoms by *NR3C1* methylation is relevant to the issue. As is evident from the SEM models, and as seen in Figures 3 and 4, there were striking similarities between the patterns of associations involving interactions between prenatal and postnatal depression and sex differences, not only for child anxious-depressed symptom but also for *NR3C1* methylation. Furthermore, in this study we reduced risks arising from multiple analyses of many potential methylation sites by examining only one site that had been identified from a meta-analysis of previous studies [24].

## CONCLUSIONS

Our findings are important in five major ways. First they provide pointers to study designs that could be introduced into animal models where mechanisms can be examined using experimentally controlled risks. These would for example examine the interplay between prenatal and postnatal risks in relation to the role of the placenta in regulating passage of maternal glucocorticoids to the foetus, which in turn can be controlled by further epigenetic modifications of specific placental genes [46]. Second they illustrate how evolutionary hypotheses regarding parental investment in offspring can be used to generate novel, and in some ways surprising, predictions regarding parenting and early development in humans. Third, testing in this way can generate further productive questions. In this study, while there was good evidence for mismatch effects in females on *NR3C1* methylation and child symptoms, and for a sex difference in relation to child symptoms, the prenatal-postnatal depression mismatch was also associated with *NR3C1* methylation in males, which was contrary to the predictions. Further study is needed into the conditions under which fetal programming effects are seen in males as well as females, and under what conditions there are sex differences in the behavioural implications of *NR3C1* methylation. Fourth they show that, even though human development is subject to many complex social and psychological influences, biological mechanisms conserved across many non-human species, can be highly influential. Fifth they suggest that some prenatal effects on epigenetic and behavioural outcomes in early childhood, differ radically in males and females, and so further study of sex specific mechanisms is needed. This will have implications for our understanding of the biology of psychiatric disorders arising in childhood.

## Abbreviations

HPA: Hypothalamic-Pituitary-Adrenal Axis
PAR: Predictive Adaptive Response
*NR3C1*: Nuclear Receptor Subfamily 3 Group C Member 1 Gene
SEM: Structural Equation Modelling
T-W: Trivers-Willard Hypothesis,
GR: Glucocorticoid Receptor
DNA: Deoxyribonucleic Acid.
CpG: Cytosine Phosphate Guanine
IMD: Index of Multiple Deprivation
EPDS: Edinburgh Postnatal Depression Scale
CBCL: Child Behavior Checklist
CNS: Central Nervous System

## DECLARATIONS

### Ethics approval and consent to participate

Ethical approval for the study was granted by the Cheshire North and West Research Ethics Committee, UK, on the 27th June 2006. The letter confirming ethical agreement for the study (reference number 05/Q1506/107) stated, ‘On behalf of the Committee, I am pleased to confirm a favourable ethical approval for the above research on the basis described in the application form, protocol and supporting document as revised.’ At recruitment at 20 weeks pregnancy, after complete description of the study to the women, written informed consent was obtained by the study midwives, who then administered questionnaires and an interview in the clinic.

### Consent for publication

Not applicable

### Availability of data and material

There is not open access to the data, because that is not permitted by our ethical approval. However requests for access to the data can be considered via contact with the first author.

### Authors' contributions

JH, HS, AP designed the study, CM, JQ conducted the methylation estimations, JH, HS, NW supervised data collection, JH, AP, NW analysed the data, JH, AP, HS, NW wrote the paper, and all authors read and approved the final manuscript.

## Acknowledgements

We are very grateful to all participating families and to the research staff who contributed to this work. We also thank Wirral University Teaching Hospital NHS Foundation Trust who supported initial recruitment and Cheshire and Wirral Partnership NHS Foundation Trust and Wirral Community NHS Trust for their current support and the NIHR Biomedical Research Centre for Mental Health at the South London and Maudsley NHS Foundation Trust and King’s College London. We would also like to extend our thanks to all the staff involved in the Wirral Child Health and Development Study.

